# Linking host plants to damage types in the fossil record of insect herbivory

**DOI:** 10.1101/2021.11.05.467393

**Authors:** Sandra R. Schachat, Jonathan L. Payne, C. Kevin Boyce

## Abstract

Studies of insect herbivory on fossilized leaves tend to focus on a few, relatively simple metrics that are agnostic to the distribution of insect damage types among host plants. More complex metrics that link particular damage types to particular host plants have the potential to address additional ecological questions, but such metrics can be biased by sampling incompleteness due to the difficulty of distinguishing the true absence of a particular interaction from the failure to detect it—a challenge that has been raised in the ecological literature. We evaluate a range of methods for characterizing the relationships between damage types and host plants by performing resampling and subsampling exercises on a variety of datasets. We found that the components of beta diversity provide a more valid, reliable, and interpretable method for comparing component communities than do bipartite network metrics. We found the rarefaction of interactions to be a valid, reliable, and interpretable method for comparing compound communities. Both beta diversity and rarefaction of interactions avoid the potential pitfalls of multiple comparisons. Lastly, we found that the host specificity of individual damage types is challenging to assess. Whereas some previously used methods are sufficiently biased by sampling incompleteness to be inappropriate for fossil herbivory data, alternatives exist that are perfectly suitable for fossil datasets with sufficient sample coverage.

## 1 Introduction

Insect herbivory on fossilized leaves (henceforth, “fossil herbivory”) has been noted incidentally for over one hundred years (Potonié, 1893). However, the systematic collection of herbivory data only came with the advent of the Damage Type system (Wilf and Labandeira, 1999), for which each type of insect damage— *e*.*g*., circular holes below 1 mm in diameter, circular holes between 1–5 mm in diameter–is assigned a unique number and is classified into a broader “functional feeding group” (Labandeira et al., 2007).

Traditionally, quantitative analyses of fossil herbivory have focused on two topics: the richness of damage type diversity at a fossil assemblage or for a particular host plant (Wilf and Labandeira, 1999), and the intensity of insect damage as measured by the percentage of leaf area removed by herbivores (Beck and Labandeira, 1998). Another layer of biological and analytical complexity can be added by linking particular host plants to particular damage types. On the one hand, quantitative methods in paleontology and ecology have progressed tremendously during the past two decades, making it possible to conduct complex analyses of fossil herbivory data with a single line of code. On the other hand, such analyses require more complete datasets than are typically available in studies of fossil herbivory. Complex analyses also rely upon far more assumptions than do traditional analyses, and as analytical complexity increases, the underlying assumptions and their effects can become more difficult to identify and address.

### 1.1 Research topics that link host plants to damage types

Three interrelated research topics link host plants to damage types: host specificity, component communities, and compound communities. Host specificity differentiates among generalist and specialist feeding strategies. A component community is the entire suite of heterotrophs that relies, directly or indirectly, on a plant taxon: its herbivores and their predators, parasitoids, and parasites (Root, 1973). A suite of coexisting component communities, *i*.*e*., those of the different plant species within the same forest, is called a “compound community” (Reice, 1974; Whittaker and Levin, 1977; Basset, 1992; Novotny et al., 2002). All of these topics present challenges when translated to the fossil record.

The host specificity of each fossil insect damage type is typically measured on a scale of 1 to 3 (Labandeira et al., 2007). Generalized damage types, occurring on a range of distantly related plant hosts, have a score of 1. Intermediate damage types have a score of 2. Specialized damage types, restricted to very closely related plant hosts, have a score of 3. These scores are assigned to damage types that occur on three or more specimens in a fossil assemblage. Damage types that occur on only one or two specimens are assigned the default score of 1, for generalized damage (Wilf and Labandeira, 1999). The assignment of these scores at various fossil assemblages is difficult to replicate because the boundaries between the scores are not defined quantitatively—the “1, 2, 3” labeling system could have used letters instead, *e*.*g*., “A, B, C”—but the many datasets that have become available since 1999 can be used for sensitivity analyses to evaluate the validity and reliability of this system.

For component communities, identification of the secondary consumers associated with the herbivores on a host plant is nearly impossible with fossils. The same is often true of the herbivores, because plants and insects are rarely preserved in meaningful quantities in the same deposits (Greenwood, 1991; Martínez-Delclòs and Martinell, 1993; Smith and Moe-Hoffman, 2007). Nonetheless, component communities in the fossil record have been widely discussed using damage types as proxies for herbivore taxa (Correia et al., 2020; Ding et al., 2014, 2015; D’Rozario et al., 2011; Feng et al., 2017; Kustatscher et al., 2018; Labandeira, 1998, 2002; Labandeira and Currano, 2013; Labandeira et al., 2013, 2016, 2018; Liu et al., 2020; Schachat et al., 2014, 2015; Slater et al., 2012, 2015; Xu et al., 2018). However, here too, there is reason for caution: even the fossil floras that have been most thoroughly sampled for insect herbivory contain various damage types that occur on only one specimen (Wilf et al., 2005, 2006; Prevec et al., 2009; Wappler, 2010; Knor et al., 2012; Wappler et al., 2012; Donovan et al., 2014; Adroit et al., 2018; Labandeira et al., 2018; Xu et al., 2018; Deng et al., 2020), indicating that many damage types remain unobserved due to incomplete sampling—and, as noted above, whether a sparsely sampled damage type is assumed under this method to be a rare generalist or a rare specialist depends on whether it was observed on two or three plant specimens. Because we cannot find every damage type from a fossil assemblage, and because we cannot link damage types to the insect taxa in a one-to-one manner, the term “component community” as developed in the context of modern ecology may be somewhat inapplicable. These issues then scale up to consideration of compound communities.

Despite these issues, the general concepts drawn from modern ecology that underlie discussions of component communities in the fossil record are nevertheless valid. Ancient plants surely had specialist and generalist herbivores that formed component communities along with their secondary consumers on each plant host species. Thus, these concepts are worthy of consideration although we must be wary of the fidelity with which those communities might be documented in the fossil record. In particular, bipartite network analysis has recently been applied to fossil herbivory datasets to address questions about host specificity and component communities (Swain et al., 2021b; Currano et al., 2021). Bipartite networks are networks that connect taxa at two trophic levels, such as plants and their herbivores or herbivores and their parasitoids. Alternatively, beta diversity (Baselga, 2010; Baselga and Orme, 2012; Baselga, 2017) and rarefaction of interactions (Dyer et al., 2010) can be used to examine herbivore specialization and component communities from the leaf damage record. Calculating the beta diversity of damage types on different host plants is a straightforward way to compare component communities. Rarefying interactions is a straightforward way to quantify the diversity of associations within a compound community. Here, these alternatives are evaluated through sensitivity analyses to determine how much sampling is required for accurate and precise results, with the aim of ascertaining whether and how quantitative methods can be used to evaluate host specificity, component communities, and compound communities in studies of fossil herbivory. Bipartite network analysis requires special consideration because of the assumptions it requires of the fossil record and because of the risks associated with the large number of metrics that are generated.

### 1.2 Theoretical issues with bipartite network analysis

#### 1.2.1 Treating damage types as analogues of herbivore taxa

Methods that link particular host plants to particular damage types often treat damage types as analogues for herbivorous insect taxa. For example, the two recent studies that performed bipartite network analysis on fossil herbivory data (Swain et al., 2021b; Currano et al., 2021) used a software package (`bipartite`; Dormann et al., 2008) intended for modern ecological networks that requires direct substitution of damage types for herbivore taxa—constituting an explicit, specific assumption that has not been substantiated and likely never can be. Only one study has used neontological data to evaluate the correlation between damage types and herbivores (Carvalho et al., 2014). In two tropical forests, the diversities of damage types and insect herbivores were found to be correlated, reaffirming the value of the traditional paleontological metric of damage type diversity. However, no claim was made as to whether the apparent specialization of a damage type reliably indicates whether the damage type was produced by a specialist herbivore.

Simple arithmetic supports the idea that specialized herbivores are responsible for many occurrences of “generalized” damage types: with hundreds of thousands of herbivorous insect species and only a few hundred damage types, no clean correspondence between insect species and damage types is possible. For example, Damage Type 012, the most common type at both forests studied by Carvalho et al. (2014), was found on all twelve host plant species examined and was caused by 50 insect species (46 of them specialists) in one locality and 37 insect species (23 of them specialists) in the other. All that complexity is collapsed into a single generalist when fossil damage types are treated as substitutes for actual herbivores. Any methods, such as bipartite network analysis, that require treating damage types as substitutes for herbivore taxa appear not to be appropriate for fossil herbivory data.

#### 1.2.2 Sampling incompleteness

All sampling of the fossil record is incomplete, but methods that link particular host plants to particular damage types are far more biased by incomplete sampling than are the methods that address the diversity and intensity of insect herbivory. For a tally of the number of insect damage types on two host plant taxa, as an example, the more completely sampled host could be iteratively subsampled down to the amount of surface area or sample coverage available for the less completely sampled host plant (Figure 1a). Although the subsampling procedure might cause a failure to detect a significant difference that would become apparent with additional sampling, any significant differences observed among the subsampled damage type diversities are likely, although not guaranteed, to reflect true differences. Thus, estimating damage type diversity by subsampling two incompletely sampled host plants is a common and uncontroversial endeavor. We do not know which specific damage types evaded detection, but we do not need to know this in order to estimate the damage type diversities of these two host plants when subsampled to the same surface area or sample coverage.

**Figure 1:**
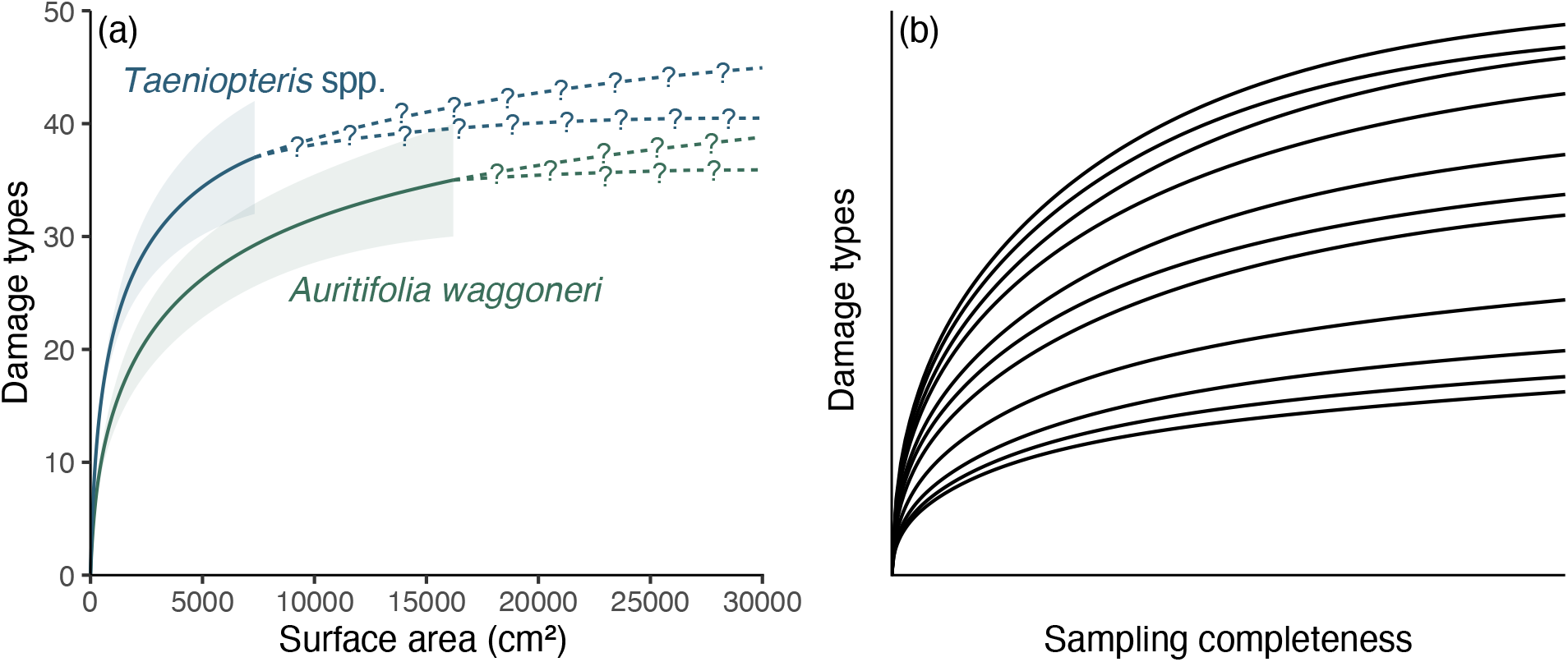
A comparison of the sampling completeness that can be expected for studies of fossil herbivory (a) with the sampling completeness needed for methods that link host plants to damage types to be unbiased by sampling completeness (b). (a) Rarefaction of damage types on the two dominant host plants at the Colwell Creek Pond assemblage. The solid lines and corresponding 84% confidence intervals represent interpolated damage type diversity, and the dashed lines with question marks represent extrapolated diversity. (b) An illustration of the sampling completeness that is needed for bipartite network analysis not to be biased by sampling: the rarefaction curve for each host plant should have sample coverage *sensu* Chao and Jost *(2012)* above 0.99. All rarefaction curves shown in this panel have coverage between 0.995 and 0.997.

When it comes to estimating host specificity or comparing component communities, however, the unknowable identities of unobserved damage types are of paramount importance. According to the criteria that have traditionally been used to assign host-specificity scores (Wilf and Labandeira, 1999), a damage type need occur on only three specimens in order to receive a host-specificity score. The data are taken at face value, and the appearance of a damage type on three leaves is deemed adequate to designate a damage type as specialized, regardless of the possibility that a fourth or fifth observation might occur on a different host and thus change the host-specificity score. The procedures used to compare component communities are incapable of distinguishing a true absence of a damage type on a host plant from the failure to detect a damage type that was present on the host. Differentiating true absences from failures to detect is known to pose tremendous difficulties in both neontological (Blasco-Moreno et al., 2019) and paleontological (Smith et al., 2021) studies.

Attempts to compare host specificity and component communities across different assemblages complicate matters even further. As an example drawn from Permian assemblages of Texas for which damage type data are available for each specimen, the amount of broadleaf area examined from Colwell Creek Pond (Schachat et al., 2014) is approximately four times that of Williamson Drive (Xu et al., 2018) and more than fifteen times that of Mitchell Creek Flats (Schachat et al., 2015) or South Ash Pasture (Maccracken and Labandeira, 2020). There is just no good way to compare host specificity and component communities across these assemblages, because subsampling Williamson Drive and Colwell Creek Pond down to the amount of surface area examined at Mitchell Creek Flats and South Ash Pasture will fundamentally change the relationships among host plants and their damage types. At Colwell Creek Pond, DT014 has been observed on two *Auritifolia waggoneri* Chaney, Mamay, DiMichele & Kerp specimens and on 20 *Taeniopteris* spp. Brongniart specimens. DT247 has been observed on 15 *A. waggoneri* specimens and 2 *Taeniopteris* spp. specimens. If the data from Colwell Creek Pond are subsampled to one-fifteenth of the original amount of surface area, the specificity coding of the damage types that are still observed at this lower level of sampling will fundamentally change: various damage types will appear more specialized than they are, and in many dimensions, the component communities of the two dominant host plants will appear more distinct than they are.

For rarefied damage type diversity and for the intensity of herbivory, the results generated at lower levels of sampling completeness are simply a less-precise, under-powered version of the results generated at higher levels of sampling completeness (Schachat et al., 2018). For component communities, however, the results generated with less sampling are fundamentally changed. In the words of Blüthgen et al. (2008), “Rarely observed species are inevitably regarded as ‘specialists,’ irrespective of their actual associations, leading to biased estimates of specialization.” Indeed, misleading results at incomplete sample sizes are exactly what biologists found when they subsampled some of the canonical datasets that have been used to construct bipartite networks (Morris et al., 2014, Figure 3) as part of the cottage industry that has emerged to evaluate how incomplete sampling biases bipartite network metrics (Goldwasser and Roughgarden, 1997; Vázquez and Aizen, 2003; Blüthgen et al., 2006, 2008; Dormann et al., 2009; Dorado et al., 2011; Gibson et al., 2011; Costa et al., 2016; Fründ et al., 2016; Jordano, 2016; Kuppler et al., 2017; Maia et al., 2018; Henriksen et al., 2019).

A related pitfall of bipartite network analysis that looms large in the neontological literature may well be insurmountable for studies of fossil herbivory: sampling evenness. Prior to the construction of bipartite networks, the sampling of fossil leaves for insect damage types should be not only complete at the level of the assemblage but should be similarly complete across all host plants within the assemblage—*i*.*e*., sampling of all host plants under consideration should be even (Gibson et al., 2011; Doré et al., 2021). In studies of modern communities, sampling evenness can be achieved in various ways, *e*.*g*., equal amounts of time being dedicated to hand-collecting of insects and equal numbers of beating samples collected for each of ten tree species (Basset et al., 1996) and equal amounts of surface area sampled for each plant species (Novotny et al., 2012). However, uniformly exhaustive sampling is a near impossibility for studies of fossil herbivory (Figure 2). Most species in a given community are rare (Diserud and Engen, 2000), and many if not most studies of fossil herbivory have examined fewer than 1,000 leaves due to a combination of small numbers of specimens preserved in the fossil record and limited time that investigators are able to invest in each study. Therefore, in studies of fossil herbivory, most plant hosts are represented by a maximum of a few hundred leaves.

**Figure 2:**
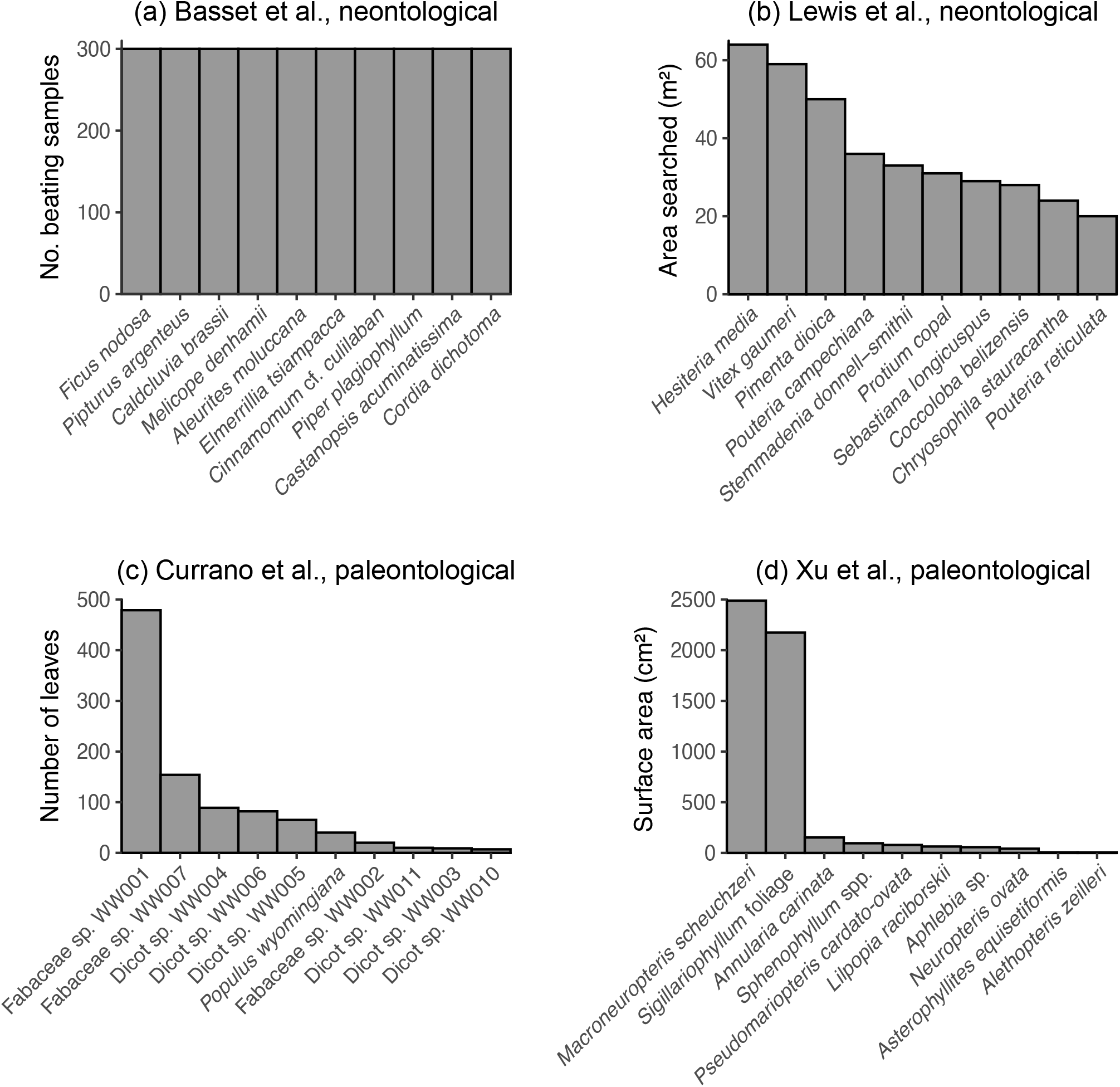
The sampling evenness for host plants in neontological (a–b) and paleontological (c–d) datasets that can be used to link host plants to herbivores or damage types. (a) Basset et al. (1996); this maximally even sampling is representative of various other neontological studies of plant–insect networks (Novotny et al., 2002, 2004, 2012; Lundgren and Olesen, 2005; Olesen et al., 2008; Pinheiro et al., 2008; Gibson et al., 2011; Grass et al., 2013; Trøjelsgaard et al., 2015; Oleques et al., 2019; Zemenick et al., 2021). (b) Lewis et al. (2002). (c) Currano et al. (2008). (d) Xu et al. (2018).

Combining the concepts of sampling completeness and evenness, Morris et al. (2014) recommended constructing bipartite networks for datasets in which all rarefaction curves—in this case, damage type diversity curves for all host plants—asymptote (Figure 1b). Various neontological food web studies have followed this recommendation (*e*.*g*., Smith-Ramírez et al., 2005; Burkle and Irwin, 2009; Mokam et al., 2014; Kemp and Ellis, 2017; Peguero et al., 2017; Bennett et al., 2018; Maia et al., 2018). However, this is not nearly as easily achieved with paleontological data as with neontological data. (Whereas one might question whether it is possible for a rarefaction curve to truly asymptote, the concept of “sample coverage” *sensu* Chao and Jost (2012) provides a measure of the slope of a rarefaction curve: when the curve has reached an asymptote, its slope equals 0 and coverage equals 1. For our purposes, sample coverage above 0.99 can be considered complete. If a dataset with ten or more host plants that have coverage above 0.99 eventually becomes available, it can be used to evaluate whether slightly lower amounts of coverage continue to yield reliable results. The Appendix lists examples of host plants that have been censused for fossil herbivory for which sample coverage of damage types is above 0.99.)

#### 1.2.3 HARKing

A “reproducibility crisis” in science (O’Boyle et al., 2017; Hutson, 2018; Nelson et al., 2021; Fraser et al., 2018; O’Dea et al., 2021; Parker et al., 2019; Bissonette, 2021) has reinforced the need for caution surrounding practices such as multiple comparisons and hypothesizing after the results are known (HARKing). In historical sciences such as paleontology, HARKing is more difficult to avoid. Understanding the properties of a large data compilation is needed to understand which analyses are feasible, but a preliminary understanding of these properties can easily lead researchers toward the questions for which a positive result is most likely.

Paleobiology cannot entirely rid itself of HARKing, but good analytical practices can identify methods that yield valid and reliable results at realistic sample sizes and that do not lend themselves to unnecessary multiple comparisons. In this context, bipartite networks present additional challenges not related to sampling. When the popular R package `bipartite` is used with its default settings to study plant–herbivore interactions, the `networklevel` function calculates 47 bipartite network metrics and the `grouplevel` function calculates 30 metrics: 15 for each host plant taxon and 15 for each herbivore taxon (Dormann et al., 2008)—77 metrics despite few studies addressing 77 distinct questions. Such a multitude of metrics raises the risk of spurious correlations whereby a small minority of metrics support preconceived notions by chance.

For bipartite network studies, calculating a single bipartite network metric per study has been recommended to avoid “metric hacking”, *i.e*., the “nonmutually exclusive use of multiple network metrics that are correlated by variables held in common [*e.g*., number of host plant taxa, or sampling completeness] and the inflation of type I error rates as a result of indiscriminate selection of network metrics, comparisons or hypotheses after analyses have been conducted” (Webber et al., 2020). However, Webber et al. (2020) also note that appropriate metric is often unclear for any given ecological question. This warning echoes concerns raised over a decade earlier: “Network analyses of mutualistic or antagonistic interactions between species are very popular, but their biological interpretations are often unclear and incautious” (Blüthgen, 2010). The unclear meanings of bipartite network metrics raise the specter of the “file drawer” problem, in which results that are inconclusive, negative, or do not fit with the authors’ agenda are not reported (Fraser et al., 2018). The complexity of bipartite networks makes their analysis subject to these risks in a way that traditional metrics of herbivore damage diversity and intensity are not.

## 2 Methods

Bipartite networks and several alternative methods were evaluated using existing data with a focus on the Willershausen assemblage (Adroit et al., 2018) as the angiosperm-dominated assemblage with a complete, publicly available dataset that has the highest number of leaves examined. Of the assemblages previously examined in the context of bipartite networks (Currano et al., 2021), Willershausen is emphasized as a conservative test because it is among the few assemblages most likely to have sufficient sampling completeness to quantify host specificity, component communities, and compound communities.

All analyses were performed with `R` version 4.1.1 (R Development Core Team, 2021). Color schemes were generated with the packages `colorbrewer` (Neuwirth and Brewer, 2014) and `scico` (Pedersen and Crameri, 2020).

### 2.1 Evaluating bipartite network analysis

#### 2.1.1 Sensitivity of bipartite network metrics to sampling completeness

The 28 network-level metrics previously named in fossil herbivory studies (Swain et al., 2021b; Currano et al., 2021; Swain et al., 2021a) that are calculated with the `networklevel` function in the `bipartite` package (Dormann et al., 2008) were calculated for the Willershausen assemblage, using subsampling and resampling procedures to evaluate their validity and reliability. Leaves that were not identified to the level of genus were removed from the dataset. Each subsampling and resampling routine was iterated 1,000 times.

In the first set of routines (“complete”), the cleaned Willershausen dataset was analyzed in its entirety, resampled to the number of leaves in the cleaned dataset (7,333), and subsampled to 3500, 1000, 500, and 300 leaves. Following previous methods (Swain et al., 2021b), all host plant taxa represented by fewer than five specimens were removed after the data were resampled or subsampled but before any analyses were performed.

In order to mirror neontological datasets (Basset et al., 1996; Lewis et al., 2002) that were recently compared to fossil herbivory data (Swain et al., 2021b), a second set of routines (“top-ten”) involved only the ten host plant taxa at Willershausen with the highest numbers of leaves, ranging from the 948 leaves of *Zelkova ungeri* Kovats down to the 164 leaves of *Betula maximowicziana* Regel. This top-ten dataset of 3602 leaves was resampled to the original number of leaves and subsampled to 1800, 1000, 500, and 300 leaves.

For the sake of comparison, we calculated damage type diversity with coverage-based rarefaction (Chao and Jost, 2012) for each resampled and subsampled dataset, using the `iNEXT` function in the `R` package `iNEXT` (Hsieh et al., 2016). We rarefied damage type diversity to the three sample coverage thresholds discussed by Schachat et al. (2021): 0.7, 0.8, and 0.9.

#### 2.1.2 Bipartite network metrics and the potential for HARKing

To evaluate the possibility of “multiple network metrics that are correlated by variables held in common”—the collinearity among metrics noted as a major pitfall of bipartite network analysis (Webber et al., 2020)—the same 28 network-level metrics discussed above were calculated for a series of fossil assemblages deposited shortly before, during, and after the Paleocene/Eocene Thermal Maximum and the Early Eocene Climatic Optimum in the Bighorn Basin and Wind River Basin. Network metrics were calculated after subsampling the data from each assemblage to 300 leaves, following the procedure of Currano et al. (2021). If a subsample is larger than 50% of the original dataset, the number of possible unique samples decreases, causing the confidence limits to narrow even though they ought to widen continuously for larger sample sizes. Therefore, subsampling to 300 leaves and generating accurate confidence intervals requires a sample size of at least 600 leaves. The ten relevant assemblages with 600 or more leaves are Skeleton Coast and Lur’d Leaves from the Bighorn Basin (Wilf et al., 2006); Dead Platypus, Daiye Spa, Hubble Bubble, the South Fork of Elk Creek, PN, and Fifteenmile Creek from the Bighorn Basin (Currano et al., 2008, 2010); and the Wind River Interior and Wind River Edge assemblages from the Wind River Basin (Currano et al., 2019).

### 2.2 Evaluating alternatives to bipartite network analysis

#### 2.2.1 Beta diversity

We evaluated the validity and reliability of measures of abundance gradients (analogous to nestedness: when the damage types observed on one host plant are a subset of the damage types observed on another host plant) and balanced variation in abundance (henceforth, “balanced variation”; analogous to turnover: when non-overlapping suites of damage types are observed on different host plants). These are the two components of beta diversity that explicitly account for differences in abundance (Baselga, 2017). Our first analysis of beta diversity focuses on the two host plants represented by the highest numbers of leaves at Willershausen: *Z. ungeri* and *Fagus sylvatica* L. We used each subsampled and resampled dataset generated from the complete Willershausen dataset. Our second analysis of beta diversity focuses on *A. waggoneri* and *Taeniopteris* spp., the two most abundant host plants at Colwell Creek Pond (Schachat et al., 2014). These two host plants were analyzed at five levels of sampling. They were jointly resampled to the original amount of surface area they comprise in the Colwell Creek Pond dataset (23,527.89 cm^2^) and were subsampled to a total of 11,750, 8,000, 4,000, and 2,000 cm^2^. Our third analysis of beta diversity focuses on *Macroneuropteris scheuchzeri* (Hoffmann) Cleal, Shute & Zodrow and foliage assigned to *Sigillariophyllum* Grand’Eury, the two most abundant host plants at Williamson Drive (Xu et al., 2018). These were jointly resampled to the original number of leaves they comprise in the Williamson Drive dataset (1524) and were subsampled to a total of 750, 600, 450, and 300 leaves. Although surface area measurements were taken for Williamson Drive (Xu et al., 2018), we subsampled these data by number of leaves because the surface area measurements for individual specimens are not available. Each subsampling routine was iterated 1,000 times.

Abundance gradients and balanced variation were calculated for each subsampled and resampled dataset using the `beta.pair.abund` function in the `R` package `betapart` (Baselga and Orme, 2012). We used the `Coverage` function in the `R` package `entropart` with the “Chao” estimator (Marcon and Hérault, 2015) to calculate sample coverage for each of the two plant hosts in each subsampling and resampling routine.

#### 2.2.2 Host specificity

The sensitivity of host specificity scores to sampling completeness was evaluated with the complete and top-ten resampling and subsampling routines for the Willershausen dataset. For each set of sampling routines, we recorded the number of host plant taxa on which we observed a randomly selected damage type within the 99th, 74th, and 49th percentiles of prevalence (Table 1).

**Table 1:** The percentiles of leaves on which damage types were observed at the Willershausen assemblage.

|  | Complete |  |  | Top-ten |  |  |
| --- | --- | --- | --- | --- | --- | --- |
|  | 99th percentile | 74th percentile | 49th percentile | 99th percentile | 74th percentile | 49th percentile |
| Number of leaves | 721 | 16 | 6 | 381 | 22 | 4 |
| Damage types | DT003 | DT033, DT145 | DT010, DT021, DT052, DT081, DT142, DT190, DT198 | DT003 | DT004, DT020 | DT008, DT052, DT061, DT168 |
| Randomly selected damage type | DT003 | DT033 | DT081 | DT003 | DT004 | DT168 |

We performed a separate sampling procedure to address the impact of absolute and relative surface area on estimates of host specificity. For this procedure we used the data from Colwell Creek Pond (Schachat et al., 2014), because this assemblage contains a large amount of surface area examined and because surface area measurements are available for each individual specimen along with damage type data. We sampled specimens belonging to *A. waggoneri, Taeniopteris* spp., *Evolsonia texana* Mamay, and *Supaia thinnfeldioides* White, with replacement, to a series of 51 equally spaced surface-area thresholds from 500 cm^2^ to 25,500 cm^2^. The smallest of these is approximately 2% of the total surface area, and the largest of these is approximately 100% of the total surface area. We resampled the data to each threshold 10,000 times, for a total of 510,000 iterations. For each iteration, we noted whether DT032 and DT120—which are distributed across all four of these host plant taxa—were restricted to only one host plant, thus falsely appearing to be specialized. If so, we noted the number of specimens on which the damage type had been observed.

#### 2.2.3 Rarefaction of interactions

The method of Dyer et al. (2010), which measures the diversity of interactions at an assemblage, can be implemented with any algorithm that performs rarefaction. We discuss considerations for coverage-based rarefaction of interactions in the Appendix.

We performed coverage-based rarefaction of interactions on data from Williamson Drive (Xu et al., 2018) and Colwell Creek Pond Schachat et al. (2014). We conducted coverage-based rarefaction on the original dataset and upon iteratively resampling each dataset to the original amount of surface area, and upon subsampling each dataset to 50% and 25% of the original surface area. (Surface area data were collected for each specimen at Williamson Drive but were not published with the damage type data. Therefore, the surface area assigned to each specimen was the mean value for the taxon to which it belongs.) We rarefied each vector of interaction counts to a sample coverage of 0.771, which is the maximum amount of coverage reached by all subsampled datasets.

To understand how rarefaction of interactions might perform on an angiosperm-dominated dataset with complete sampling, we simulated a vector of counts of interactions using the base-`R` function `rlnorm` with the settings `meanlog=0` and `sdlog=1.5`. This procedure generated 3,000 values, which we had to round to whole integers because these values represent simulated counts. Upon removing the values that round down to 0, we had 2,046 simulated unique interactions which had were observed a total of 9,597 times. These numbers are approximately double those seen in the Willershausen dataset, so we attributed these simulated interactions to 15,000 leaves because this is approximately double the number in the Willershausen dataset.

We examined the validity and reliability of rarefaction of interactions in this simulated dataset by subsampling. We subsampled the interactions to one half of the original count (4,798), attributing these to one half of the original number of leaves (7,500). We then subsampled the interactions to one quarter of the original count (2,399), attributing these to one half of the original number of leaves (3,750). We rarefied each vector of subsampled interaction counts to a sample coverage of 0.726, which is the maximum amount of coverage reached by all subsampled datasets.

All rarefaction of interactions was carried out with the `estimateD` function in the `R` package `iNEXT`. All resampling and subsampling procedures were iterated 1,000 times.

## 3 Results and Discussion

### 3.1 Sensitivity of bipartite network metrics to sampling completeness

None of the 28 network-level metrics mentioned in previous studies of fossil herbivory (Swain et al., 2021b,a; Currano et al., 2021) perform as unbiased estimators for the complete Willershausen dataset (Figure 3). (An unbiased estimator is an estimator whose average value does not change in response to sampling completeness.) Two simple criteria for robustness to sampling completeness are that the 95% confidence intervals for all subsampling routines contain the mean estimate for the resampling routine, and the 95% confidence interval for the resampling routine contains the mean estimates for all subsampling routines. Coverage-based rarefaction of damage type diversity fulfills these two criteria (Figure 4), but not a single network metric examined here does.

**Figure 3:**
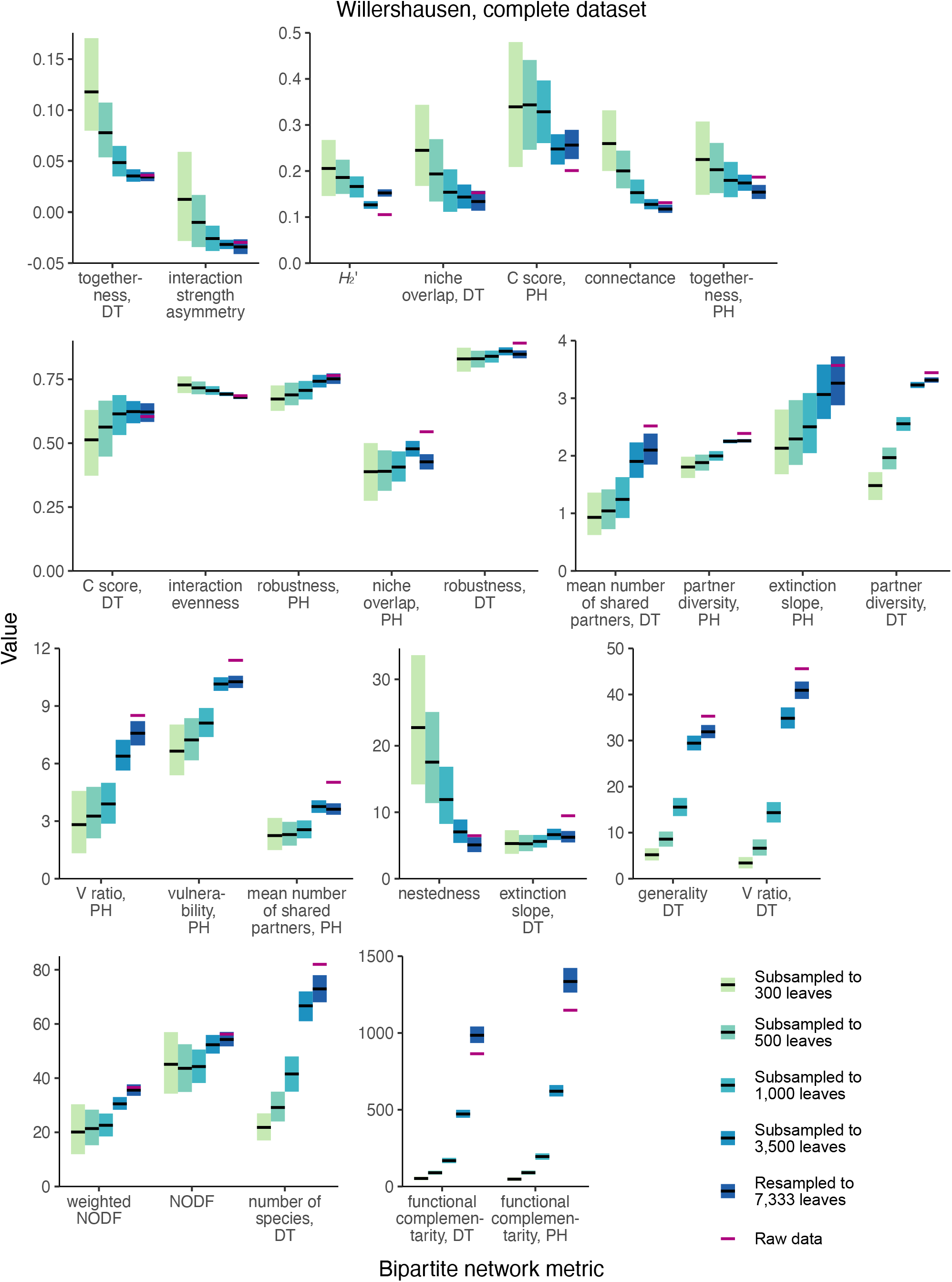
Mean values and 95% confidence intervals for bipartite network metrics, generated by resampling and subsampling the cleaned Willershausen dataset in its entirety. Legend: PH = plant host, DT = damage type.

**Figure 4:**
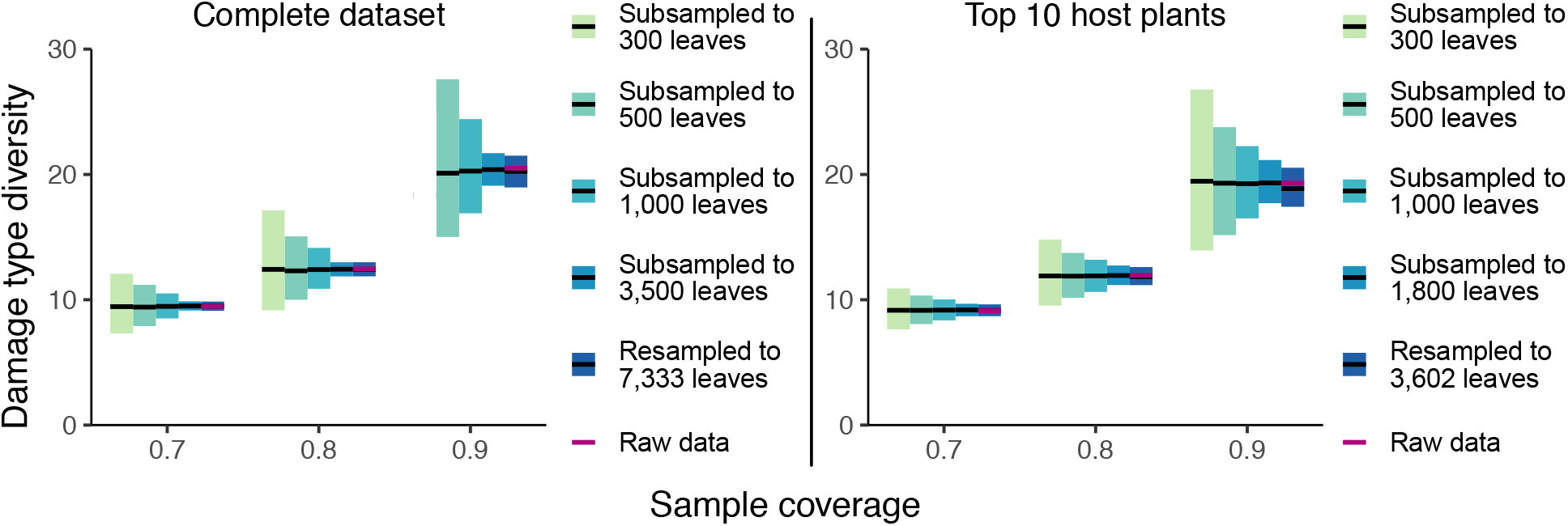
An example of a nearly unbiased estimator. Mean values and 95% confidence intervals for coverage-based rarefaction, generated by resampling and subsampling the Willershausen dataset. Moreover, coverage-based rarefaction performs as a consistent estimator, in that estimates converge on the true value as sample size increases. No results are presented for 300 subsampled leaves from the complete dataset at sample coverage of 0.9 because some iterations of this sampling routine yielded an observed sample coverage below 0.9.

When the Willershausen data are restricted to only the ten host plants with the highest number of leaves in the dataset (Figure 5), 4/28 network metrics fulfill these criteria and thus perform comparably well to coverage-based rarefaction: togetherness for damage types, niche overlap for damage types, C score for damage types, and nestedness.

**Figure 5:**
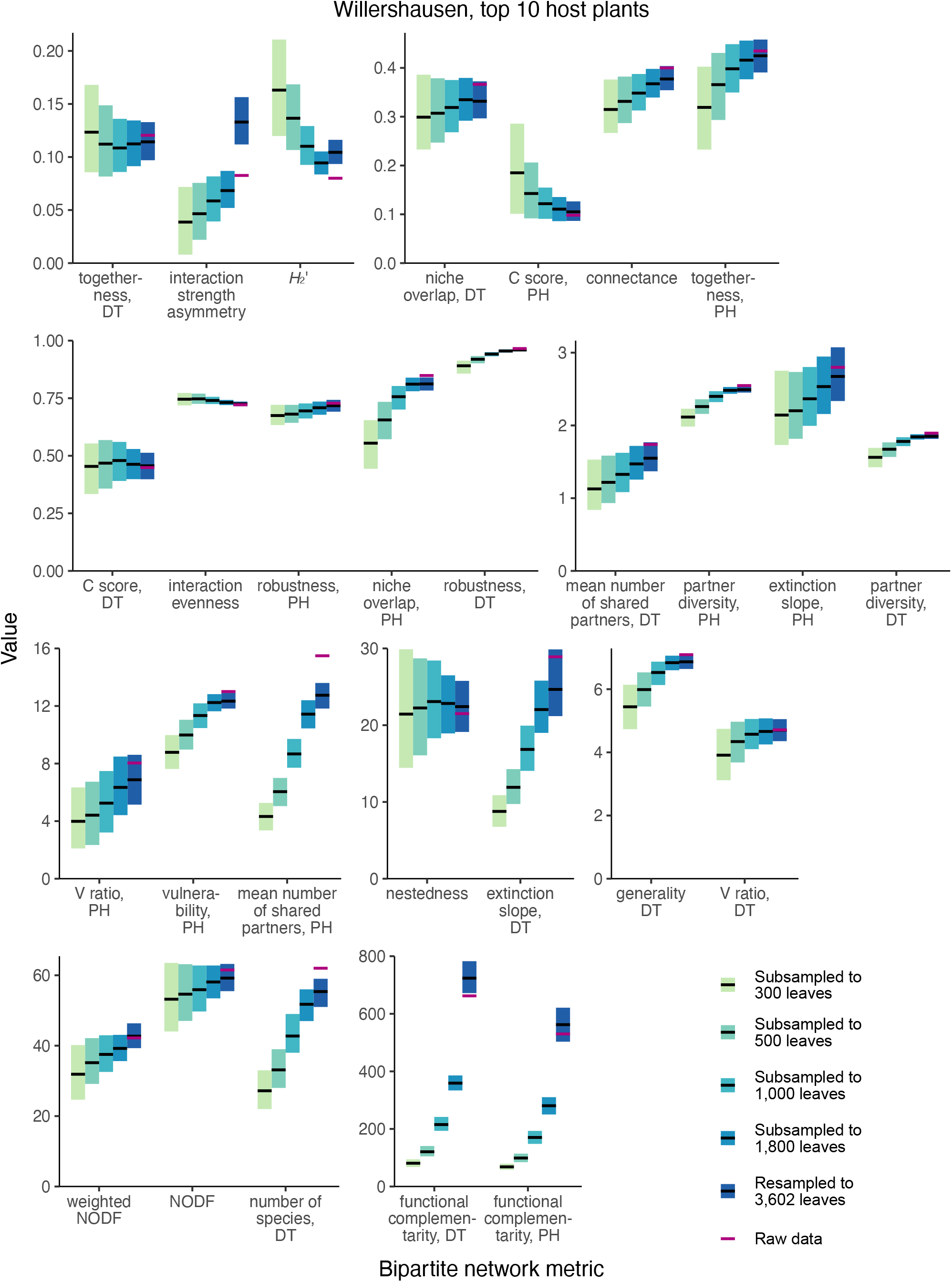
Mean values and 95% confidence intervals for bipartite network metrics, generated by resampling and subsampling data for the ten host plants at Willershausen represented by the highest numbers of leaves. Legend: PH = plant host, DT = damage type.

Only one network metric, C score for damage types, is among the best-performing in both the complete and top-ten analyses of the Willershausen dataset. If the C score for damage types were found to be robust for the majority of available fossil herbivory datasets, which are far less complete than Willershausen, a key question would still need to be answered: What does the C score tell us? Many metrics are generated with little understanding of what they mean in practice, simply because their calculation requires only a few lines of code. The C score has been described in the fossil herbivory literature as “the checkerboard (mutual presence/absence) nature of the interactions” (Swain et al., 2021b) and as “the randomness of species distribution across an ecosystem” (Currano et al., 2021), but no outstanding paleontological questions that can be addressed with such a metric have been identified.

#### 3.1.1 Apparent robustness at lower sample sizes

For many metrics in both the complete and top-ten datasets, the mean estimate and the limits of the confidence intervals change little for the subsampling routines at 1,000, 500, and 300 leaves. However, when the resampling routine and the subsampling routines with over 1,000 leaves are taken into account, it is clear that these metrics are biased by sampling incompleteness. The misleading, apparent lack of bias in certain network metrics seen at lower levels of sampling makes intuitive sense. When a relatively large proportion of realized interactions are unobserved because only 1,000 leaves have been sampled, the additional proportion of realized interactions that go unobserved at 500 or 300 leaves will make little difference for various metrics. These findings and this reasoning highlight the danger of evaluating the bias of network metrics by performing sensitivity analyses on smaller datasets. Therefore, any metrics that appear robust to subsampling routines performed on datasets smaller than that of Willershausen should be treated with extreme caution. For these same reasons, methods that quantify the extent to which bipartite network metrics are biased by sampling incompleteness (Swain et al., 2021a) may well be unreliable, especially when applied to incomplete datasets.

#### 3.1.2 Implications for other assemblages

At any amount of sampling that is realistic for studies of fossil herbivory, the results of bipartite network analysis are biased by sampling completeness. The finding that certain metrics are “relatively robust” (Swain et al., 2021a) is an inevitability by chance alone given presentation of dozens of metrics (Swain et al., 2021b; Currano et al., 2021). Even when we limit our analysis to the ten most abundant host plants at Willershausen, the mean estimates at 300 and 500 leaves for the best-performing metrics (Swain et al., 2021a) either lie beyond (NODF, *H* _2_*’*, connectance, and niche overlap PH) or just barely fall within (niche overlap DT) the 95% confidence interval generated with the resampled dataset. Estimates of these metrics at different sampling intensities are even more discordant for the complete Willershausen dataset.

Neontological evaluations of bipartite networks have indicated that sampling is complete enough for bipartite network metrics to be valid and reliable only when two criteria are met. First, the rarefaction curves should asymptote for all taxa at the lower trophic level (Arceo-Gómez et al., 2018), *e.g*., rarefaction curves of damage type diversity for each host plant under consideration in a study of fossil herbivory, should reach sample coverage above 0.99. At Willershausen, coverage for the top ten host plants ranges from 0.90 to 0.99. However, at Castle Rock (Wilf et al., 2006), another of the few assemblages with over 2,000 angiosperm leaves examined for which damage type data are available for each specimen, coverage of the top ten host plants is much lower, with some taxa preserving no damage at all and the highest coverage only reaching 0.72. At the Bílina–DSH assemblage (Knor et al., 2012), also with over 2,000 angiosperm leaves examined, coverage of the top ten host plants ranges from 0.59 to 0.90. Therefore, low sample coverage of damage types for individual host plants is clearly not due to lack of investigator effort; this is a characteristic of some of the best-sampled assemblages. Rather, low sample coverage of damage types for individual host plants is a near-inevitability given the vastly uneven frequencies of both host plants and damage types in fossil assemblages. Even the less common host plants must be represented by enough specimens for their individual damage diversity rarefaction curves to asymptote. This requirement is unrealistic for essentially the entirety of the fossil record as it is currently sampled.

### 3.2 Alternatives to bipartite networks

#### 3.2.1 Beta diversity

Our calculations of balanced turnover and abundance gradients for the two dominant host plants at Willershausen show that these metrics are valid and reliable under the resampling routine and under the routine in which the dataset was subsampled to 3,500 leaves (Figure 6). At lower levels of sampling, the abundance gradient metric remains valid but is noticeably less reliable. The balanced variation metric becomes less valid and reliable at lower levels of sampling. Unsurprisingly, estimates of balanced turnover and abundance gradients are most valid and reliable when coverage is high.

**Figure 6:**
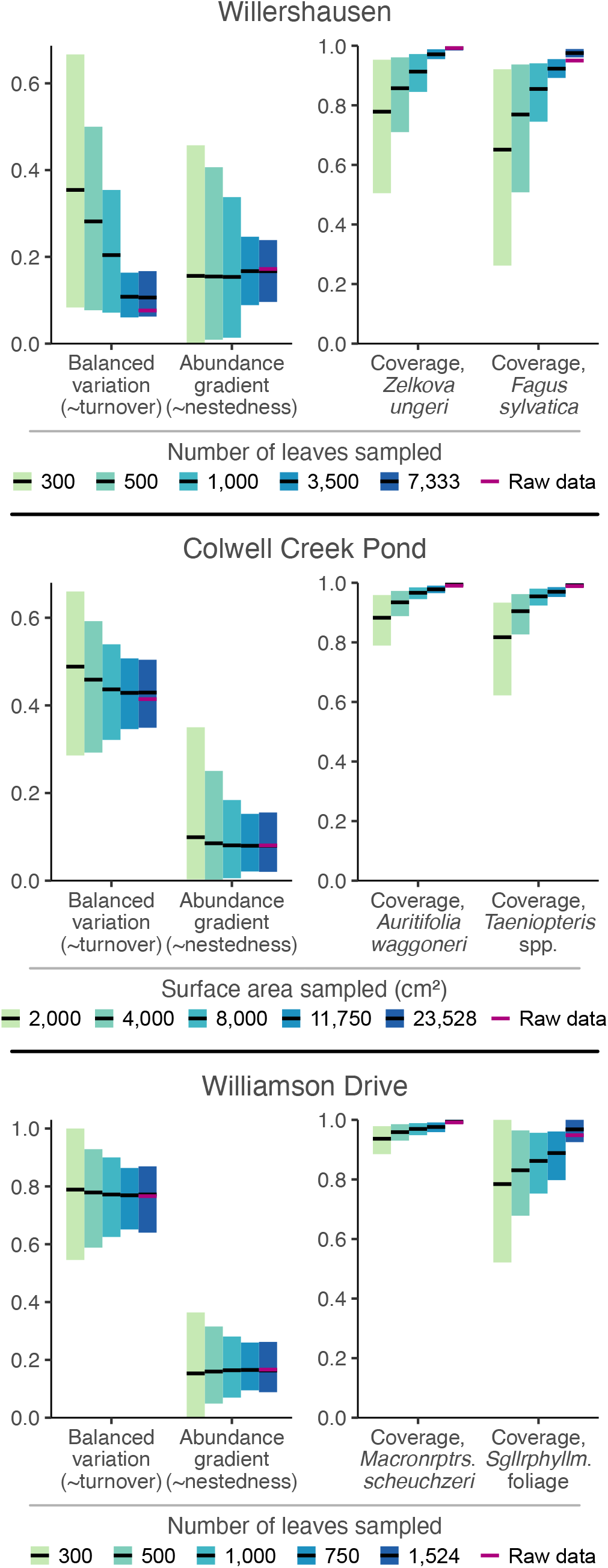
Mean values and 95% confidence intervals for beta diversity metrics, generated by resampling and subsampling data for the two most abundant host plants from Willershausen, Colwell Creek Pond, and Williamson Drive.

Among the datasets generated by iteratively resampling the Willershausen data and by subsampling the data to 3,500 leaves, coverage estimates do not overlap but estimates of balanced turnover and abundance gradients overlap almost perfectly. However, estimates become much less reliable when the Willershausen dataset is subsampled to only 1,000 leaves, and the levels of coverage for *Z. ungeri* and *F. sylvatica* fall to 0.91 and 0.86, respectively.

The Colwell Creek Pond data yield much more valid and reliable results. This is perhaps unsurprising, because coverage of the second-most abundant host plant is higher at Colwell Creek Pond than at Willershausen. Whereas it is very rare for two host plants within a single assemblage to have such high sample coverage—0.990 for *A. waggoneri*, and 0.989 for *Taeniopteris* spp.—our findings suggest that valid and reliable estimates of balanced turnover and abundance gradients are achievable for those rare assemblages with two host plants that are nearly completely sampled.

The Williamson Drive data yield results that are even more valid and reliable than those for Colwell Creek Pond. This is a bit surprising: although the most dominant host plant at Williamson Drive, *Macroneuropteris scheuchzeri*, has sample coverage of 0.991, the second-most dominant host plant, *Sigillariophyllum* foliage, has sample coverage of only 0.948—far less than that of *Taeniopteris* spp. at Colwell Creek Pond. For Williamson Drive, balanced variation and abundance gradients essentially perform as unbiased and consistent estimators, to nearly the same extent as does coverage-based rarefaction (Figure 4). Further analyses are needed to determine exactly why these two metrics perform somewhat better for the Paleozoic data than for Willershausen—richness of damage types may be a key determinant—and particularly why these metrics perform better for Williamson Drive than for Colwell Creek Pond.

Nevertheless, it is clear that these two components of beta diversity are a preferable alternative to bipartite network metrics. They are more valid and reliable than nearly any bipartite network metric that has been examined for fossil herbivory (Currano et al., 2021; Swain et al., 2021b). Their meanings are clear, as is the difference between them. They provide no opportunity for metric hacking. They can be calculated for pairwise comparisons among host plants, or can be used to generate a single value for an entire assemblage (Baselga and Orme, 2012; Baselga, 2017), and can thus be used whether an assemblage contains two or twenty host plants with nearly complete sampling.

#### 3.2.2 Host specificity

The results of our resampling and subsampling procedures demonstrate that the traditional method for assigning host specificity scores is strongly biased by sampling completeness: at lower levels of sampling, the host breadth of a damage type inevitably decreases (Figure 7). For example, in the Colwell Creek Pond resampling routines, we treated each iteration in which the generalist DT032 or DT120 damage type was restricted to only one host plant taxon as a false positive finding of specialization. DT032 appeared on only one host plant taxon in 2.72% of iterations; DT120, 3.78%. When a finding of specialization requires a damage type to appear on three or more specimens, following the convention established by Wilf and Labandeira (1999), the false positive rate falls to 0.93% for DT032 but remains at 3.34% for DT120.

**Figure 7:**
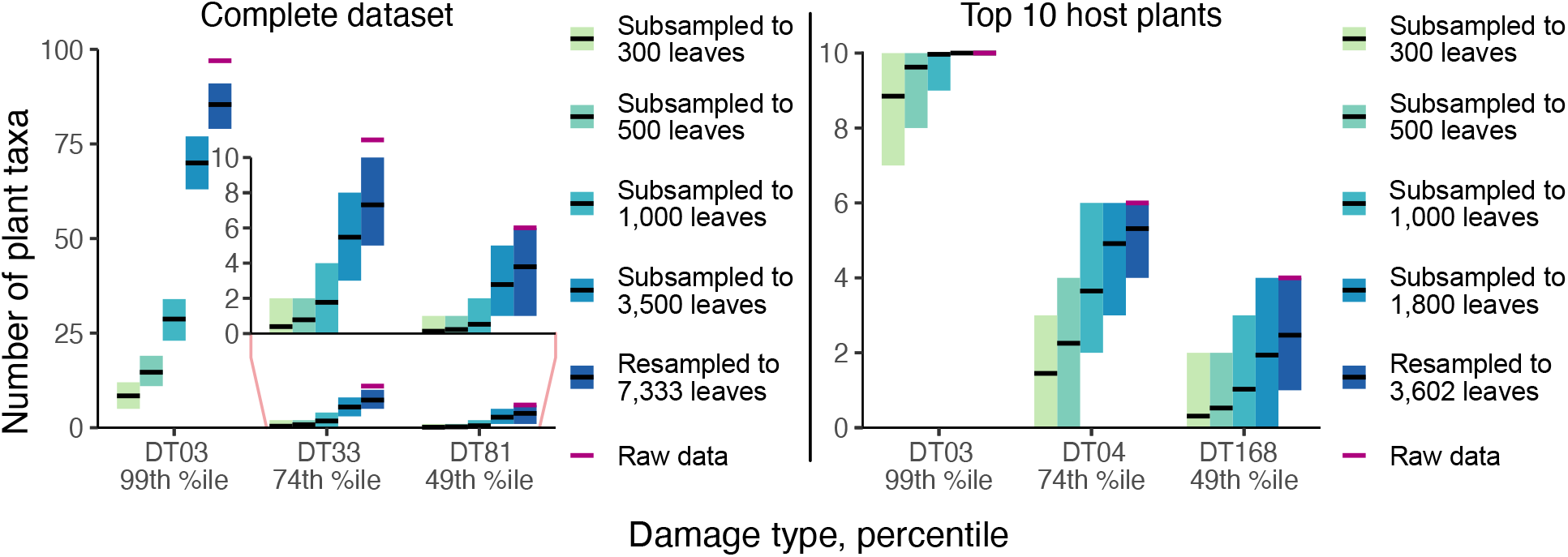
Mean values and 95% confidence intervals for the number of plant taxa on which various damage types appear, calculated with the Willershausen dataset.

The inadequacy of the three-specimen threshold for designation of a damage type as “specialized” is shown by the frequencies of false positive results (Figure 8). These frequencies follow lognormal distributions. For DT032, which was observed on fewer leaves than DT120, *σ >* 1 such that the greatest proportion of false positive results occur when this damage type is observed on only one specimen. However, for DT120, *σ <* 1 such that 4.7% of false positive results occur when this damage type is observed on only one specimen, 8.7% occur when this damage type is observed on four specimens, and 4.9% occur when this damage type is observed on nine specimens. Thus, the three-specimen threshold protects against only a small fraction of false positives.

**Figure 8:**
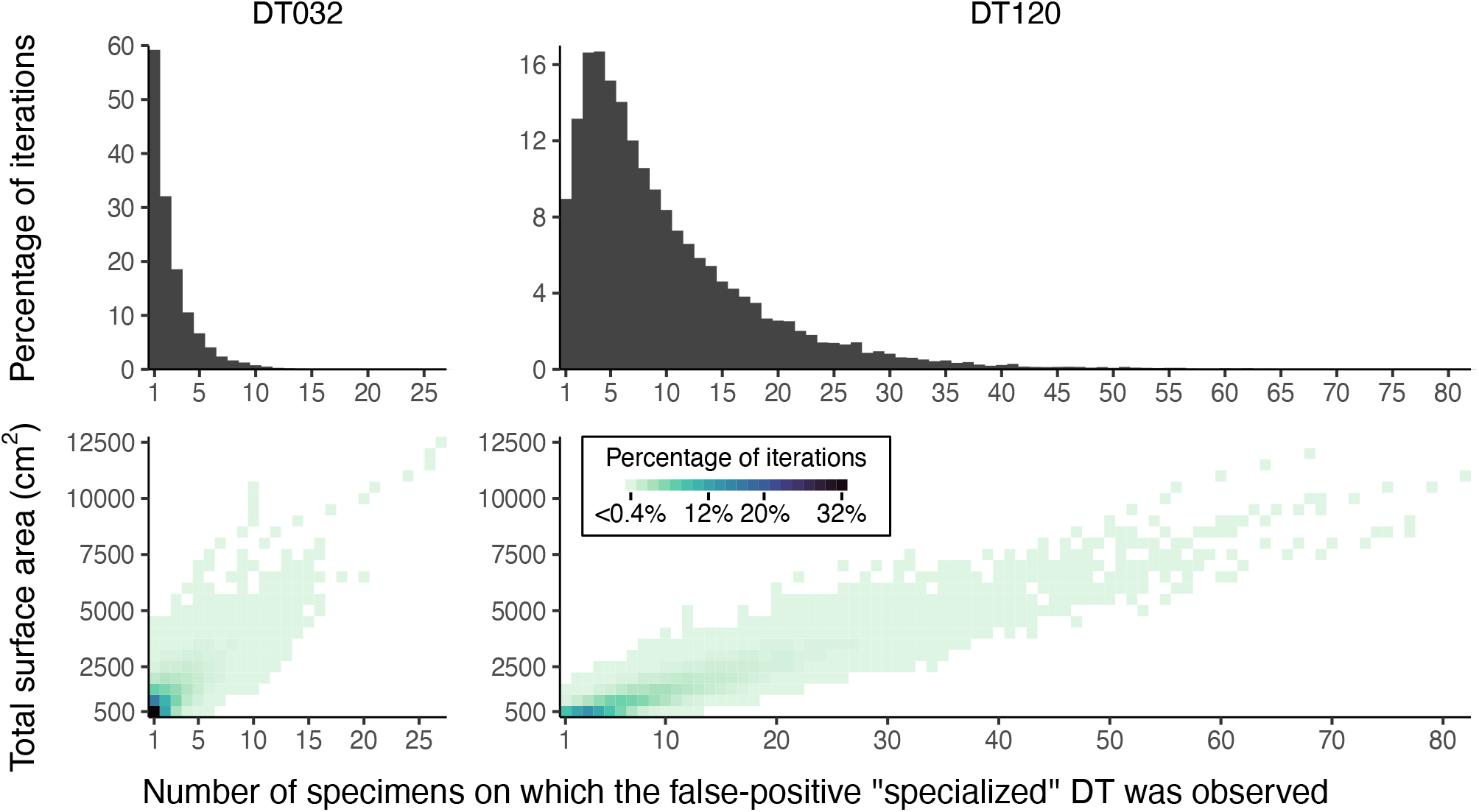
False positive results of “specialized” damage generated by iteratively resampling data from Colwell Creek Pond. We treated each iteration in which DT032 or DT120 was observed on only one host plant taxon as a false positive. The heatmaps show the percentage of iterations for each amount of subsampled surface area in which a false positive result was recovered, arranged by the number of specimens on which the damage type was observed. The histograms show the summed percentages, by number of specimens.

#### 3.2.3 Rarefaction of interactions

Coverage-based rarefaction of interactions performs as an unbiased and consistent estimator: as sampling completeness decreases, the mean estimate changes negligibly while confidence intervals widen (Figure 9). Resampled estimates and confidence intervals are often invalid for rarefaction of interactions, because the number of singletons in a resampled dataset tends not to exceed the number of singletons in the original dataset. The number of singletons is one of the main determinants of estimated sample coverage, and thus, resampled datasets tend to have higher estimated coverage than the original datasets. This means that coverage-based rarefaction will generate lower estimates for resampled data than for subsampled data. This is abundantly clear for rarefaction of interactions in the simulated dataset and is also quite notable for Williamson Drive. The estimation of confidence limits from iteratively sampled data should therefore be performed with subsampled, rather than resampled, data whenever the mean estimate generated with resampled data is clearly invalid. The methodology of coverage-based rarefaction of interactions is illustrated in Figure 10.

**Figure 9:**
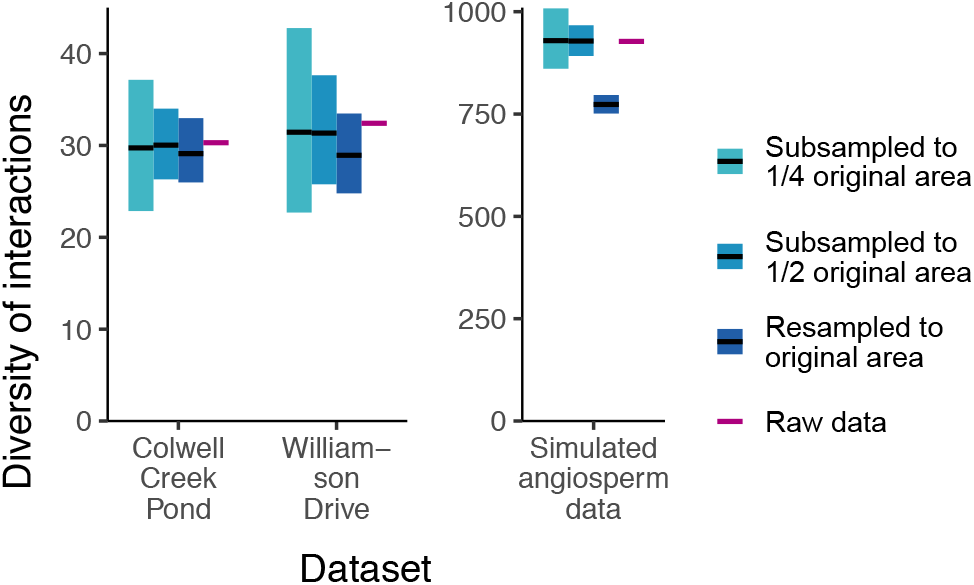
Mean values and 95% confidence intervals for coverage-based rarefaction of interactions. The datasets presented here are Williamson Drive and Colwell Creek Pond, both from the Permian of Texas (rarefied to a sample coverage of 0.771), and a simulated dataset that mimics the patterns seen among angiosperms at Willershausen (rarefied to a sample coverage of 0.726).

**Figure 10:**
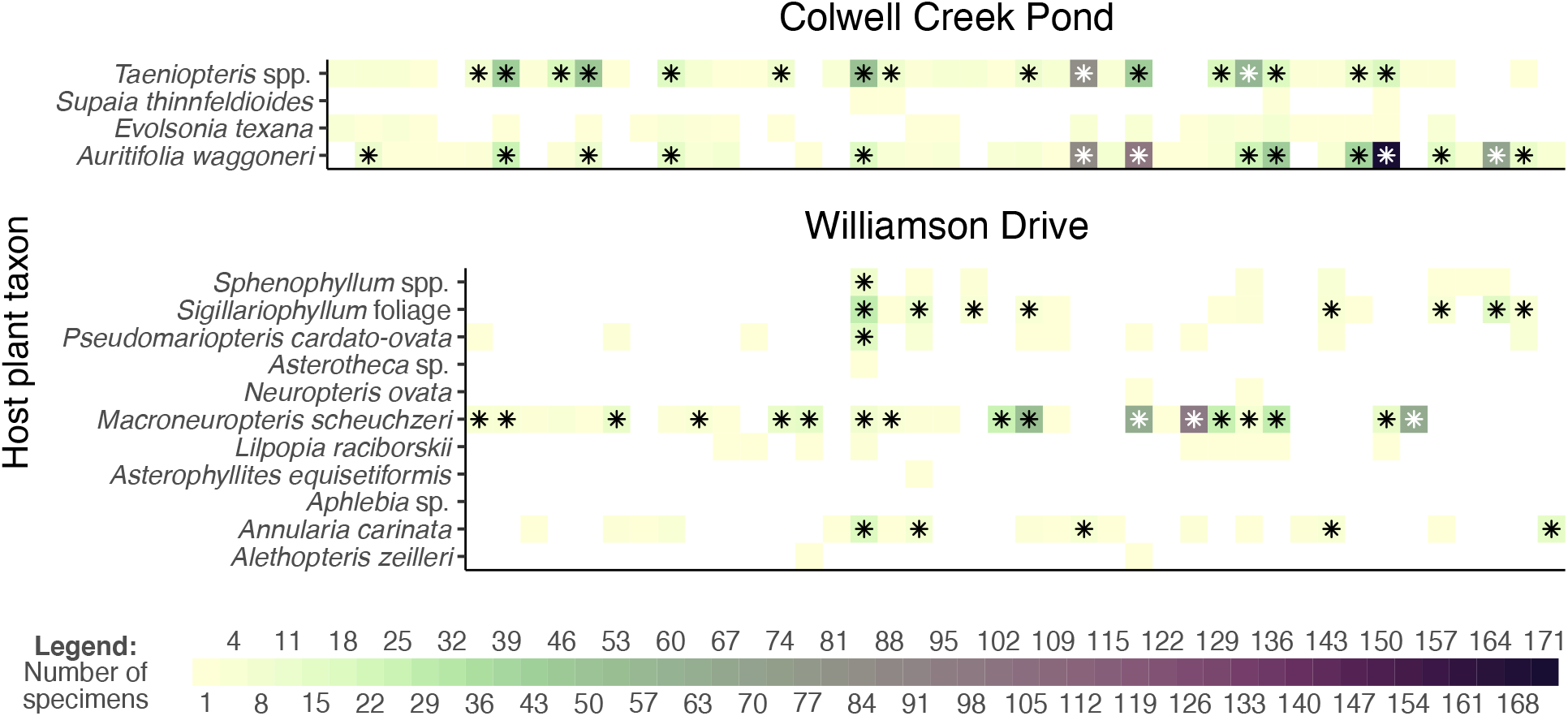
Comparison of the raw and rarefied interaction data from Colwell Creek Pond and Williamson Drive. Each column of each graph represents a damage type. The heatmaps show the prevalence of each interaction, and the asterisks denote interactions that remain after rarefying data from each assemblage to a sample coverage of 0.771.

### 3.3 An example of bipartite network metrics and the potential for metric hacking

While it has been argued that bipartite network metrics allow a more finely resolved, “in-depth” understanding of the relationships between host plants and damage types (Swain et al., 2021a), others argue that the multiple comparisons presented in many network studies often contain spurious results (Webber et al., 2020). To evaluate which of these two views of multiple comparisons in network studies is applicable to fossil herbivory datasets, we calculated bipartite network metrics for one of the most iconic and intensely studied series of assemblages in this discipline: Paleocene and Eocene floras of the western interior of North America (Wilf and Labandeira, 1999; Currano et al., 2008, 2010). The finding of increased insect herbivory at the Paleocene/Eocene Thermal Maximum (PETM) is supported by quantitative measures of herbivorized leaf area (Currano et al., 2016) and by damage type diversity, whether rarefied by number of leaves (Currano et al., 2010)—an older practice shown to be biased by differences in leaf surface area among host plant taxa (Schachat et al., 2018)—or rarefied by sample coverage (Schachat et al., 2021). Changes in herbivory at the Early Eocene Climatic Optimum (EECO) have not been examined as thoroughly (Currano et al., 2019), but the logic about climate, nutrient availability, and herbivory used to describe the PETM (Currano et al., 2008, 2010) ought to apply to the EECO as well.

When the 28 bipartite network metrics considered here are calculated for the Paleocene–Eocene assemblages of the Bighorn Basin and Wind River basin (Figure 11), none of these metrics yield extreme values for the PETM Hubble Bubble assemblage (Currano et al., 2008) or the EECO Wind River Interior assemblage (Currano et al., 2019). If these metrics are taken at face value, rather than being dismissed due to their susceptibility to sampling bias, the metrics suggest that extreme climate change does not have a perceptible impact on plant–insect interactions. For a variety of metrics (interaction strength asymmetry, the C score for host plants, connectance, togetherness, partner diversity for damage types, generality for damage types), it not the assemblage deposited during the PETM, but the assemblage deposited just afterward, that yields the most extreme values. This assemblage, South Fork of Elk Creek, was immediately noted for having only two host plants preserved in meaningful quantities (Currano et al., 2008; Currano, 2009): a peculiarity that has not been ascribed with ecological significance (Currano et al., 2008; Currano, 2009; Currano et al., 2010). However, this long-known peculiarity appears to be driving temporal patterns in approximately one quarter of bipartite network metrics. (For all other assemblages shown in Figure 11, the mean number of host plant taxa in each subsampling iteration ranges from 4.7 to 11.9.)

**Figure 11:**
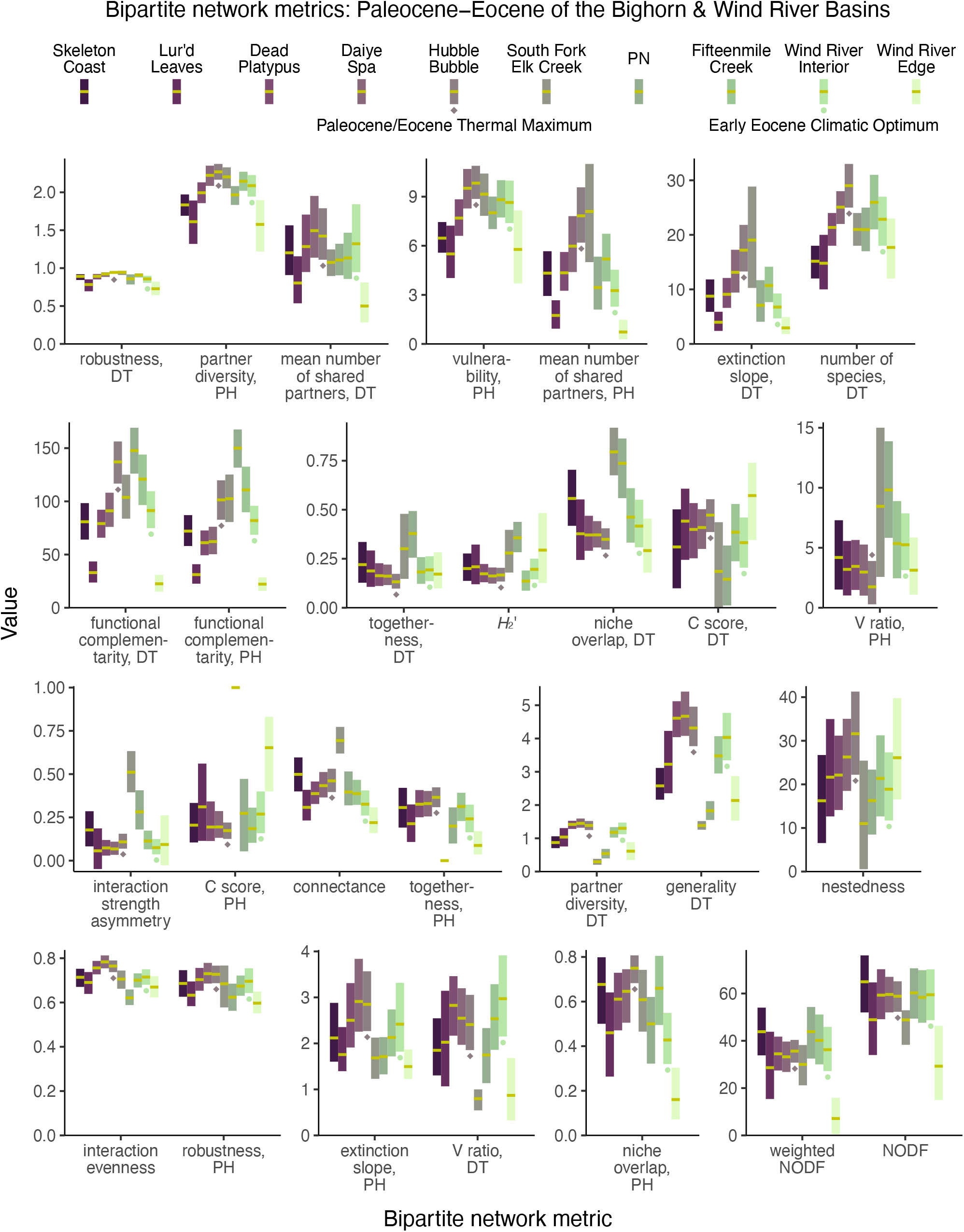
Mean values and 95% confidence intervals for bipartite network metrics, generated by subsampling each dataset to 300 leaves. Legend: PH = plant host, DT = damage type.

Different combinations of these metrics support different narratives. Of the 28 bipartite network metrics, approximately one third suggest that the PETM and EECO had similar impacts on the relationship between host plants and damage types, approximately one third suggest that the PETM and EECO had similar impacts, and approximately one third yield inconclusive results (Figure 11, Table 2). The PETM itself yields a variety of possible conclusions. Over two thirds of these metrics suggest that the relationship between host plants and damage types did not drastically change from the very late Paleocene to the PETM, and less than one quarter are inconclusive (Figure 11, Table 2). The only two metrics that suggest a drastic change in the relationship between host plants and damage types at the PETM—functional complementarity for host plants, and for damage types—are the two metrics that show the greatest amount of spread overall (Figures 3, 5, 11). Moving from the PETM into the Eocene, more than one quarter of these metrics suggest that the relationship between host plants and damage types did not change from the PETM to its immediate aftermath, over one third suggest that this relationship did indeed change, and over a quarter are inconclusive (Figure 11, Table 2).

**Table 2:**
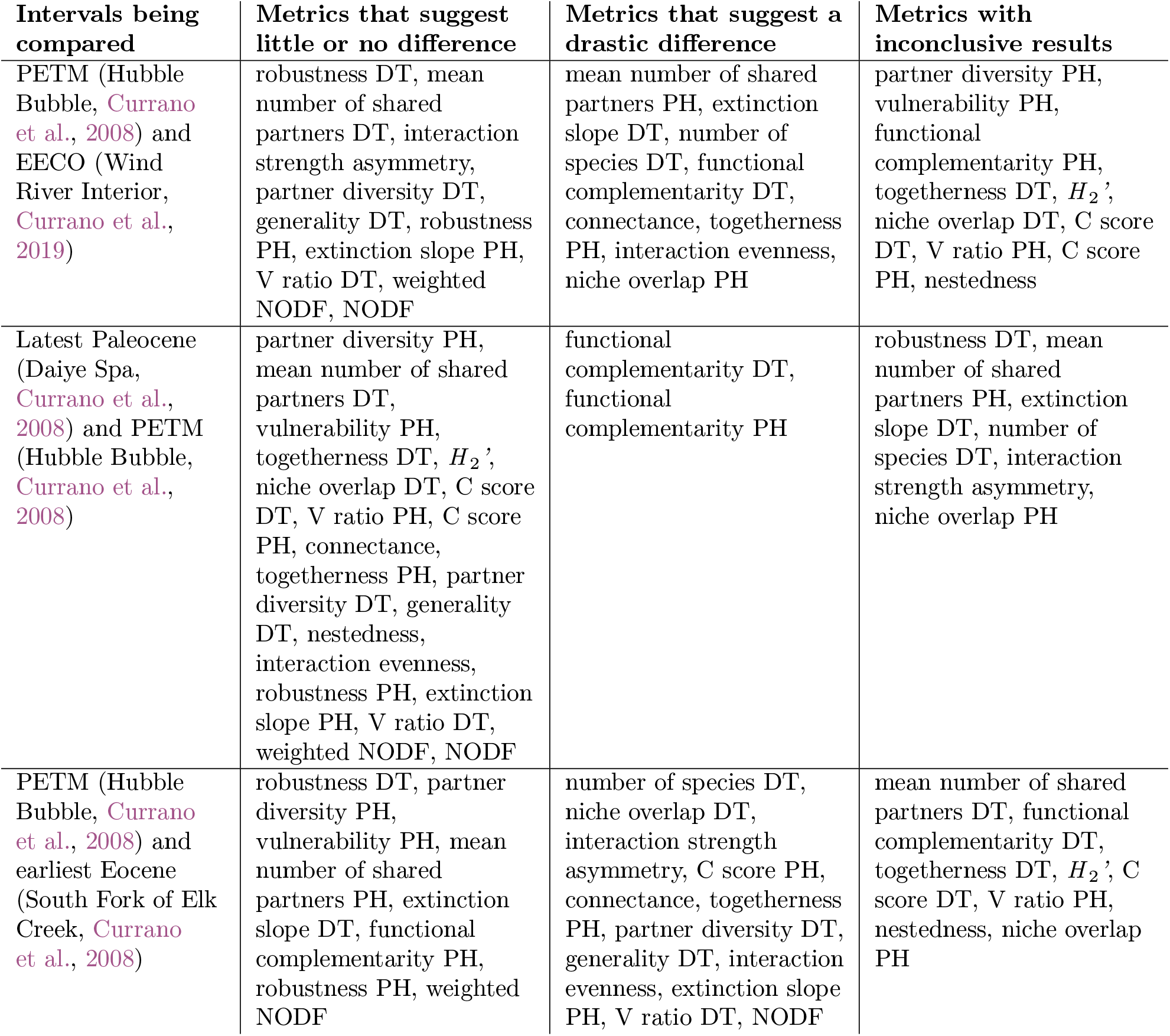
The variety of narratives about the PETM supported by different combinations of bipartite network metrics.

The only metric that returns a more extreme value for the PETM than for the two assemblages that immediately predate and postdate it—*i.e*., the mean value for the PETM lies beyond the 95% confidence intervals for any of these four other assemblages—is “number of species, DT.” We have presented this metric here as if it were a bipartite network metric, because it was previously reported as such (Swain et al., 2021b; Currano et al., 2021) and because it is calculated with the `networklevel` function in the `bipartite` package in `R` (Dormann et al., 2008). However, this is not truly a bipartite network property in that it does not respond to the distribution of damage types among the host plants.

Bipartite network properties fail to identify the PETM as an anomaly. This finding necessitates a reckoning as to whether bipartite network analysis provides additional nuance and context to traditional metrics such as the herbivory index and rarefied damage type diversity, or, alternatively, whether these metrics are too biased at realistic sample sizes to provide results that warrant interpretation. If the canonical notion of uniquely intense and diverse insect herbivory at the PETM is erroneous, that notion should of course be challenged. But, for the many reasons detailed above, the various narratives that emerge from bipartite network analysis that contradict the accepted influence of the PETM on insect herbivory are quite likely artifacts of sampling incompleteness and unevenness.

## 4 Conclusions

The challenge of linking host plants to damage types through bipartite network analysis is twofold. First, sampling incompleteness does not simply cause increased uncertainty, as is the case for consistent and unbiased estimators such as the herbivory index or coverage-based rarefaction of damage type diversity; instead, sampling incompleteness typically leads to inaccurate, misleading results. And second, the wide variety of bipartite network metrics creates many opportunities for HARKing. Those opportunities are exacerbated by the unclear meanings of these metrics.

No amount of sampling completeness can remove the potential for HARKing presented by bipartite network analysis, but our results show that alternative methods that are insusceptible to HARKing can be used to evaluate host specificity, to compare component communities, and to measure the diversity of interactions at an assemblage. Rarefied interaction richness and the components of beta diversity are much more likely than bipartite network metrics to perform as unbiased and consistent estimators, and do not require complete sampling of damage types across all host plants at an assemblage. Much essential information is still lacking: the exact sample coverage required for valid measurement of abundance gradients, balanced variation, and the diversity of interactions; as well as the surface area data required for evaluation of host specificity, which are unavailable for most published assemblages. However, the first step is understanding which analyses are meaningful and which measurements are needed for those analyses to be valid.

At present, there are a number of large gaps in our knowledge of fossil herbivory. First is the nearly complete lack of Pennsylvanian or Jurassic assemblages examined for herbivory and the lack of early-to-mid Cretaceous assemblages. Second is the general lack of assemblages examined from tropical latitudes. Third is the widespread lack of surface area measurements, which are necessary for evaluating the intensity of herbivory (Schachat et al., 2018). Fourth is the widespread lack of counts of the number of times that each damage type appears on each leaf. These data can be used to evaluate various hypotheses about the causes of increased herbivory (Schachat et al., 2021). In light of the limited amount of time that paleontologists are able to spend collecting fossil herbivory data, we believe that addressing these four gaps is the most important use of investigator effort.

## 5 Appendix

### 5.1 Calculating *p*-values for host specificity

The absolute amount of surface area examined should be taken into account when determining host specificity because if the total amount of surface area is very small, the apparent restriction of a damage type to a particular clade of host plants will very possibly be an artifact of insufficient sampling. The relative amount of surface area should be taken into account because this determines the probability that a damage type would falsely appear to be restricted to a particular clade of host plants.

Consider a hypothetical assemblage in which 100,000 cm^2^ of surface area have been examined. If DT001 is restricted to a clade of host plants represented by a mere 500 cm^2^ of surface area, and if DT001 is found on all 15 specimens belonging to the clade at this assemblage, then DT001 indeed appears to be specialized. This finding is supported by the large amount of surface area examined, by the moderately high number of specimens on which DT001 has been found, and by the small amount of relative surface area belonging to the plant clade in question, which confers a low probability that all detected incidents of DT001 would be restricted to this clade due to chance alone.

However, at Colwell Creek Pond, the host plant *Auritifolia waggoneri* accounts for over 60% of the broadleaf surface area examined. Therefore, especially if the total amount of surface area examined is low, a generalized damage type may appear to be restricted to *A. waggoneri* due to chance alone—particularly if the damage type is observed on only a few specimens. To test the frequency with which this sort of false positive finding of specialized herbivory may occur, we resampled the data from Colwell Creek Pond for the four host plant taxa from this assemblage that unambiguously meet the criteria for inclusion outlined by Swain et al. (2021b): *A. waggoneri* (63% of total broadleaf surface area), *Taeniopteris* spp. (28%), *Evolsonia texana* (9%), and *Supaia thinnfeldioides* (1%). Our analysis focuses on two damage types, DT032 and DT120. Both of these damage types occur on all four of these host plants, with distributions that approximate the amount of surface area examined for each host plant: the majority of incidences of each damage type are on *A. waggoneri* (63–89%), followed by *Taeniopteris* spp. (10–25%), *E. texana* (1–10%), and, lastly, *S. thinnfeldioides* (1–3%).

When a damage type is observed only on one clade of host plants at an assemblage, the surface area of those host plants can be used to test the null hypothesis that the damage type is restricted to a certain plant clade simply by chance. The proportion of all surface area examined at the assemblage that belongs to the clade in question—whether it is a genus or species, implying specialized host specificity, or a higher clade implying intermediate specificity—can be raised to the number of specimens on which the damage type was observed. This process generates a *p*-value that can be used to test the null hypothesis of generalized host specificity. Consider an example in which a damage type appears to have an intermediate host specificity because it occurs only on plants belonging to the same order. If this order accounts for 40% of all surface area examined at the assemblage, and if the damage type has been observed on five specimens, the *p*-value for its host specificity is 0.4^5^ = 0.01024. This value is below 0.05, and thus, the damage type has been observed on enough specimens to reject the null hypothesis of generalized host specificity. However, a correction for multiple comparisons, such as the Bonferroni correction or the Benjamini–Hochberg correction, should be used if this procedure is carried out for more than one damage type.

These findings presented in our Results section suggest that the more conservative Bonferroni correction should be used instead of the Benjamini–Hochberg correction when host specificity *p*-values are calculated for multiple damage types. Surface area data from additional assemblages, with as much area as Colwell Creek Pond or more, are needed in order to determine whether the Benjamini–Hochberg correction will suffice.

Another fundamental, unresolved issue pertaining to the assignment of host-specificity scores is the definition of “specialized” and “intermediate” host specialization. If a damage type occurs on multiple genera within the same family, is it a specialized damage type, because it is restricted to one family, or is it an intermediate damage type, because it occurs on multiple genera? To our knowledge, this question has never been answered, leaving each team of authors to draw the boundaries between specialized, intermediate, and generalized host specificity wherever they please. To our knowledge, the locations of these boundaries are not typically articulated in publications, leading to a lack of reproducibility. Because the majority of herbivorous insects feed on plants belonging to a single family (Forister et al., 2015), we recommend that a damage type which occurs on a single family be considered “specialized” and that a damage type which occurs on multiple families within a single order be considered “intermediate.”

We do not advocate assigning host-specificity scores to damage types. For reasons outlined in the Introduction, specialist herbivores can be largely or entirely responsible for a “generalized” damage type. For reasons outlined in the Results and Discussion, a “generalized” damage type can appear to be “specialized” due to sampling incompleteness. However, should any research teams continue to assign host-specificity scores, our method for generating *p*-values protects against false positive findings of specialized herbivory and our recommended boundaries between specialized, intermediate, and generalized host specificity provide an objective, reproducible, working definition.

### 5.2 Considerations for coverage-based rarefaction of interactions

The input used for bipartite network analysis and for rarefaction of interactions is essentially the same (Table 3). Bipartite network analysis uses a matrix in which each row represents a host plant, each column represents an herbivore (or, for fossil herbivory, a damage type), and each cell represents the number of times that a given interaction was observed. In the example shown in Table 3, DT001 was observed on one specimen belonging to plant sp. 1 and DT002 was observed on five specimens belonging to plant sp. 1. For rarefaction of interactions, the matrix is vectorized, or transformed into a single row. The information about particular host plants and damage types is removed, only the numbers of observations remain, the ordering of these observations does not matter, and it does not matter whether unobserved interactions with a value of 0 are retained in the vector.

**Table 3:** A toy example of the input used for bipartite network analysis. For rarefaction of interactions (Dyer et al., 2010), the input would be a vectorized version of this matrix, which could take any of the following forms: [1 5 0 2 2 0 0 6 0 0 1 0 0 1 0 1 0 3 0 1], or [1 5 2 2 6 1 1 1 3 1], or [6 5 3 2 2 1 1 1 1 1 0 0 0 0 0 0 0 0 0 0], or [6 5 3 2 2 1 1 1 1 1].

|  | DT001 | DT002 | DT003 | DT004 |
| --- | --- | --- | --- | --- |
| Plant sp. 1 | 1 | 5 | 0 | 2 |
| Plant sp. 2 | 2 | 0 | 0 | 6 |
| Plant sp. 3 | 0 | 0 | 1 | 0 |
| Plant sp. 4 | 0 | 1 | 0 | 1 |
| Plant sp. 5 | 0 | 3 | 0 | 1 |

This vector is then used for a subsampling procedure, and can be subsampled to a threshold of sample coverage as Schachat et al. (2021) have advocated. Whereas bipartite network analysis produces misleading results with incomplete sampling by treating rare, undetected interactions as true absences, rarefaction of interactions subsamples the observed interactions such that the rare, undetected interactions are removed from the dataset and thus cannot bias the results. Once the dataset for an assemblage reaches the coverage threshold to which all assemblages are subsampled, additional sampling completeness—revisiting an assemblage that already reaches a sample coverage of 0.9, and collecting additional data until sample coverage reaches 0.95—will not change the results on average, in contrast to bipartite network analysis. This is because the progression of an unbiased sampling routine will lead to additional observations of common interactions while allowing the observation of new, rare interactions.

In a typical rarefaction analysis in the context of fossil herbivory, the input is a vector that contains the number of specimens upon which each damage type has been observed. For example, if DT001 and DT002 have each been observed on three specimens and DT003 has been observed on one specimen, the input vector would take the form of [3 3 1]. To rarefy the interactions rather than the damage type incidences in this toy example, if DT001 was observed on three specimens belonging to the same plant host and DT002 was observed on two different plant hosts, the input vector would take the form of [3 2 1 1]: the second 3 in the original vector, corresponding to DT002, has been split into a 2, representing two incidences of this damage type on one plant host, and a 1, representing an incidence of this same damage type on a different plant host.

There is a computational issue with increasing the number of values in an input vector that equal 1: this reduces sample coverage (Good, 1953). Because scaling rarefaction curves by the number of leaves examined is an inadequate substitute for scaling by the amount of surface area examined (Schachat et al., 2018), coverage-based rarefaction is the only appropriate method for comparing assemblages that lack measurements of surface area. But the sampling completeness that is needed to rarefy damage type diversity (Schachat et al., 2021) will far fall short of the sampling completeness needed to rarefy the diversity of interactions. For example, when we iteratively subsampled the Willershausen dataset to 1,000 leaves, sample coverage was as low as 0.599—a level at which comparisons will be grossly under-powered, as discussed by (Schachat et al., 2021). Therefore, we evaluated rarefaction of interactions with a simulated dataset.

### 5.3 Host plants with sample coverage above 0.99

The following is a non-exhaustive list of host plants censused for fossil herbivory, for which sample coverage is above 0.99. *Zelkova ungeri* from Willershausen (Adroit et al., 2018); *Macginitiea gracilis* (Lesquereux) Wolfe & Wehr from PN (Currano et al., 2010); *Heidiphyllum elongatum* (Morris) Retallack from Aasvoëlberg 311 (Labandeira et al., 2018); *Sphenobaiera schenckii* (Feistmantel) Florin from Birds River 111 (Labandeira et al., 2018); *Platanus raynoldsii* Newberry from Mexican Hat (Wilf et al., 2006; Donovan et al., 2014); *Macroneuropteris scheuchzeri* from Williamson Drive (Xu et al., 2018); *A. waggoneri* from Colwell Creek Pond (Schachat et al., 2014); *Quercus* sp. L. from Longmen (Su et al., 2015).

When coverage equals 1, this is typically misleading, as it most likely signifies that either no damage has been found on the host plant taxon in question (the `Coverage` function in the `entropart` package calculates coverage of 1 when there is no damage) or that the sample size is much too small, which can spuriously lead to no singleton damage types. For example, Fabaceae sp. WW042 at the PN assemblage (Currano et al., 2010) is represented by 16 leaves. Three damage types are observed: DT002 is on two leaves, DT012 is on six leaves, and DT032 is on two leaves. Coverage equals 1. However, if the number of leaves with DT002 is experimentally reduced from two to one, coverage falls from 1 to 0.9111. The only host plant we are aware of for which coverage of 1 is not a spurious artifact is *Quercus* sp. from Longmen (Su et al., 2015). Twelve damage types were found on the 1,027 leaves examined. All damage types were found on at least five leaves. If the number of leaf specimens with DT045 is experimentally reduced from five to four, coverage remains at 1. This suggests a rule of thumb for determining whether a high coverage estimate is an artifact: if coverage remains above 0.99 after one leaf specimen with the rarest non-singleton damage type is experimentally removed from the dataset, the coverage estimate is indeed robust. Notably, when we subsampled the Willershausen dataset to at least 1,000 leaves and iterated this procedure 10,000 times, coverage never exceeded 0.9972. It therefore appears that all coverage estimates that equal 1 would become slightly lower—if not far lower—with additional sampling. Thus, a coverage estimate of 0.995 is a stronger indicator of complete sampling than is a coverage estimate of 1.

